# TransposonUltimate: software for transposon classification, annotation and detection

**DOI:** 10.1101/2021.04.30.442214

**Authors:** Kevin Riehl, Cristian Riccio, Eric A. Miska, Martin Hemberg

**Author notes:** Equal contributor.

## Abstract

**Motivation:** Most genomes harbor a large number of transposons, and they play an important role in evolution and gene regulation. They are also of interest to clinicians as they are involved in several diseases, including cancer and neurodegeneration. Although several methods for transposon identification are available, they are often highly specialised towards specific tasks or classes of transposons, and they lack common standards such as a unified taxonomy scheme and output file format. Moreover, many methods are difficult to install, poorly documented, and difficult to reproduce.

**Results:** We present TransposonUltimate, a powerful bundle of three modules for transposon classification, annotation, and detection of transposition events. TransposonUltimate comes as a Conda package under the GPL-3.0 licence, is well documented and it is easy to install. We benchmark the classification module on the large *TransposonDB* covering over 891,051 sequences to demonstrate that it outperforms the currently best existing solutions. The annotation and detection modules combine sixteen existing softwares, and we illustrate its use by annotating *Caenorhabditis elegans*, *Rhizophagus irregularis* and *Oryza sativa subs. japonica* genomes. Finally, we use the detection module to discover 29,554 transposition events in the genomes of twenty wild type strains of *Caenorhabditis elegans*.

**Availability:** Running software and source code available on https://github.com/DerKevinRiehl/TransposonClassifierRFSB. Databases, assemblies, annotations and further findings can be downloaded from https://cellgeni.cog.sanger.ac.uk/browser.html?shared=transposonultimate.

## 1 Introduction

Transposons are evolutionary ancient mobile genetic elements that can move via copy&paste and cut&paste transposition mechanisms. They can be classified within a taxonomic scheme (Fig. 1A), and each class is associated with a set of characteristics, e.g. proteins relevant for transposition and structural features (Fig. 1 B). During transposition, transposable elements (TEs) can leave structural patterns both at the insertion and the deletion site [1, 2, 3]. Autonomous transposons encode the tools necessary for transposition events, e.g. genes producing transposase, integrase and other enzymes [3], while non-autonomous transposons depend on proteins encoded elsewhere [4]. As the insertion of a transposon can be detrimental, many species have developed repression mechanisms, e.g. TE promoter methylation [5] and piRNAs [6]. Even though transposition events occur rarely [7], in many organisms large sections of DNA consist of either transposons or their transposition-incompetent descendants that have accumulated mutations over time [4]. It is estimated that transposons make up a large share of the genome in many species; 45% in humans, 20% in fruit flies, 40% in mice, 77% in frogs and 85% in maize [8].

**Figure 1.**
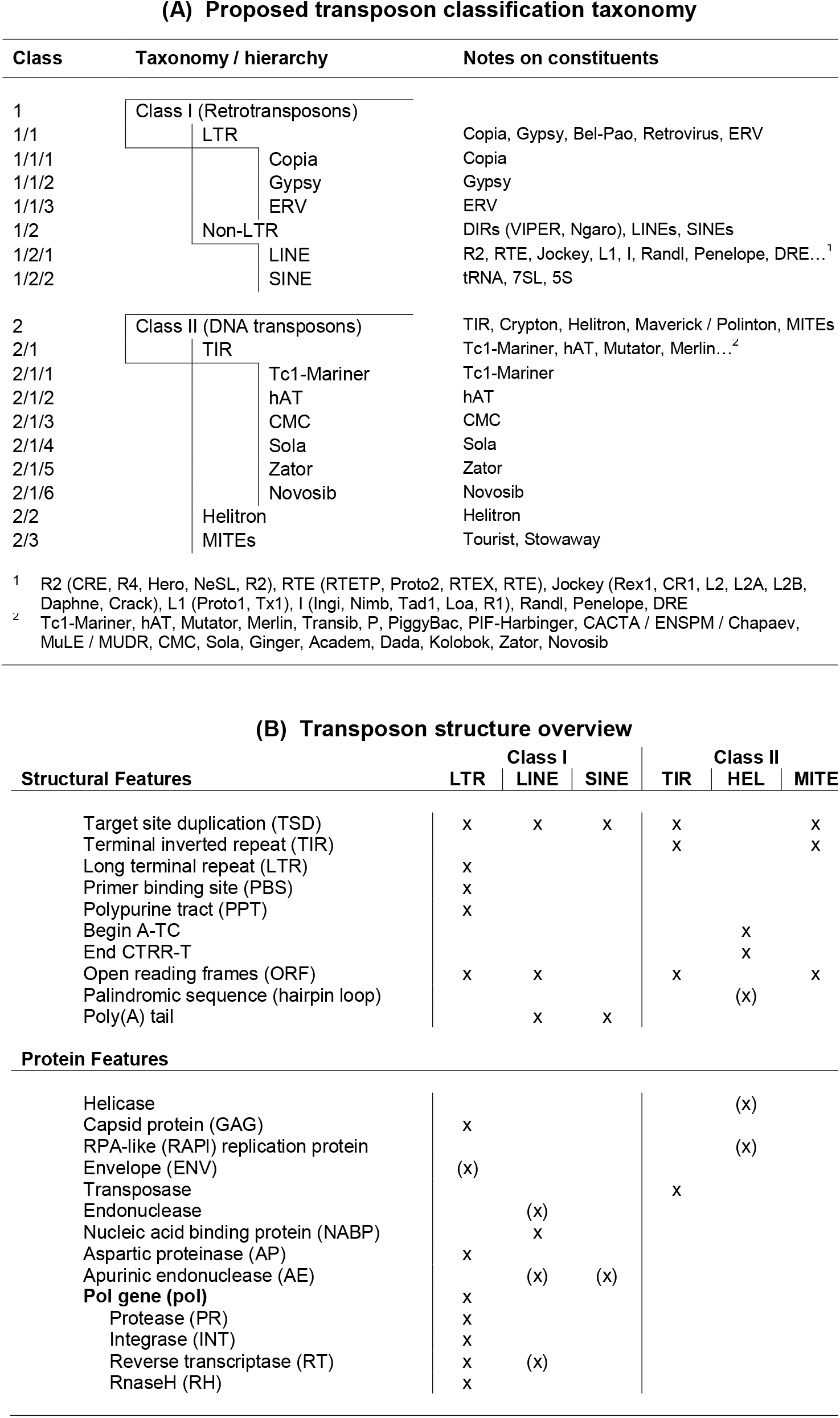
Transposon taxonomy and transposon structure. (A) The taxonomy used in this study is based on multiple classification schemes [49], [36], [106], [3] and the taxonomies used by the transposon databases. (B) Autonomous, transposition competent transposons have characteristic structural and protein features depending on their class. The proteins are necessary for the transposons to move via class-specific transposition mechanisms. The x mark which structural and protein features are characteristic to different transposon classes and sub classes for complete, autonomous transposons. The (x) mark features that are not required but if present are indicative.

Studying TEs is highly relevant for understanding evolutionary processes [9], developmental biology, gene regulation, and many diseases called transposonpathies such as subtypes of haemophilia, immunodeficiency, cancer and Alzheimer’s disease [10, 11, 12]. Also, TEs are popular for genetic engineering purposes as they allow for direct insertion of their genetic cargo into a target genome [13, 14, 15]. However, the repetitive nature of transposons and their descendants is a challenge for their analysis and discovery, in particular when using short-read sequencing technologies [7]. Long-read technologies facilitate studies of transposons and their functional consequences, but they also require novel computational tools. Although various approaches for identifying transposons have been proposed recently [16], current tools are error prone, not robust, mostly rely on prior knowledge of transposon sequences, and are often limited to a family of transposons or a group of species [17].

Here, we present a bundle of tools addressing three different tasks related to transposon identification: classification, annotation and detection. The goal of classification is to determine which taxonomic class a given transposon sequence belongs to. The annotation task consists of scanning a genome sequence to mark all transposons. Finally, the detection task involves the comparison of two genomes to identify structural variants arising from the insertion of TEs.

Existing transposon classifiers are difficult to compare directly since they vary in their approach, which features and taxonomies they use, how they evaluate predictions, and which databases are used for training. Applications of SVMs [18], hidden Markov models [19], random forests [20], Gaussian naive Bayes [21], decision trees [22], stacking [23, 24], boosting [25, 26], neural networks [27, 28, 29], evolutionary algorithms [30, 21] and genetic algorithms [31, 32, 33, 34] can be found in the literature. Most methods use sequence features, such as the k-mer frequency, the occurrence of structural [35] and protein features [18] for classification. Besides, another approach is to classify TEs using the similarity to known transposons based on a sequence library [36].

The annotation of transposons in nucleotide sequences is challenging due to the presence of transposition-incompetent TEs that have been mutated, truncated, degraded, fragmented and dismembered due to nesting [37]. Annotation is further complicated by a lack of standards [38] and disagreement on definition, taxonomy and terminology [39, 40]. Since transposons do not adhere to a universal structure [41], many researchers have employed class-specific approaches [42]. Moreover, most of the software employed for transposon annotation was originally designed for gene annotation, neglecting the peculiarities of transposons [39]. Existing transposon annotation methods (Table 1) can be assigned to one or more approaches [43, 2, 1, 41]. The *de novo* approach finds transposons by identifying repetitive sequences. It is effective in discovering previously unknown transposons with high prevalence [41], but it is computationally costly [41, 39], unable to find degraded transposons [41], and risks misidentifying repetitive DNA or high copy number genes as transposons [44, 45]. The structure-based approach (also called motif-based [42] or signature-based approach [2]) is based on knowledge of the structure of transposons and annotates by finding combinations of characteristic patterns [38, 46]. This approach enables the discovery of transposition-incompetent transposons thanks to their unique structural properties [41]. However, these approaches are often characterised by high false discovery rates [37, 44] and they miss transposons with weak signatures [37]. The similarity-based approach (also called library-based approach [2]) employs a library of known transposons together with BLAST(-like) tools. The high accuracy [41] and short runtimes [44, 47] of this approach come at the cost of its inability to find unrelated transposons [41, 47] and the dependency on quality and exhaustiveness of the library [38, 44, 48]. Moreover, the current version of the most widely used database RepBase [49] is behind a paywall and the related tools RepeatMasker and RepeatModeler are not transparent with regards to how transposons were curated and consensus sequences were generated [39].

**Table 1.** Overview of common transposon annotation tools. The most commonly used tools such as RepeatMasker and RepeatModeler cover a variety of transposons, while others focus on certain classes only. The tools use one or more of the *de novo*, structural and similarity-based transposon annotation approaches.

| Name | Approach |  |  | Class I |  |  | Class II |  |  |
| --- | --- | --- | --- | --- | --- | --- | --- | --- | --- |
|  | Novo. | Struc. | Simil. | LTR | LINE | SINE | TIR | HEL | MITE |
| RepeatMasker | x |  | x | x | x | x | x | x | x |
| RepeatModeler | x |  |  | x | x | x | x | x | x |
| CLARI-TE [107] | x | x | x | x | x | x | x | x | x |
| TESeeker [41] |  |  | x | x | x | x | x | x | x |
| PILER [40] | x |  |  | x | x | x | x | x | x |
| Censor [108] | x |  |  | x | x | x | x | x | x |
| RepLong [109] | x |  |  | x | x | x | x | x | x |
| EDTA [44] | x | x | x | x | x | x | x | x | x |
| MGEScan [110] | x | x | x | x | x | x |  |  |  |
| LTR-Finder [111] |  | x |  | x |  |  |  |  |  |
| LtrDetector [112] |  | x |  | x |  |  |  |  |  |
| LTRpred [73] | x | x | x | x |  |  |  |  |  |
| LTRharvest [66] | x | x | x | x |  |  |  |  |  |
| LTRdigest [113] |  | x |  | x |  |  |  |  |  |
| SINE-Finder [68] | x | x |  |  |  | x |  |  |  |
| SINE-Scan [69] | x | x |  |  |  | x |  |  |  |
| TIRvish [67] |  | x |  |  |  |  | x |  |  |
| HelitronScanner [42] |  | x |  |  |  |  |  | x |  |
| MUSTv2 [70] |  | x |  |  |  |  |  |  | x |
| MiteFinderII [71] |  | x |  |  |  |  |  |  | x |
| MITE-Tracker [72] |  | x |  |  |  |  |  |  | x |
| detectMITE [45] |  | x |  |  |  |  |  |  | x |
| MITE-Hunter [47] |  | x |  |  |  |  |  |  | x |

Previous efforts to detect transposition events by comparing two genomes have been based on the analysis of the depth of coverage, discordant and split read pairs [50, 51]. However, both the task of detecting structural variants (SVs) and annotating TEs are very challenging when using short reads [7]. Recently, long-reads technologies have become more widely available, but to the best of our knowledge the only existing method that can take advantage of them for TE detection is LoRTE [52]. Although results indicate that LoRTE performs well even on low coverage reads, it is limited to PacBio data and insertion and deletion SVs only.

Here, we present TransposonUltimate, a set of tools for the identification of transposons, consisting of three modules for accurate classification, annotation in nucleotide sequences and detection of transposition events (Fig. 2). Our new classifier is benchmarked against existing softwares, and we use the annotation module to analyse the genomes of three different species. Finally, the detection module is employed to identify transposition events in 20 high quality genomes from *Caenorhabditis elegans* wild isolates that were assembled using a combination of long- and short-read technologies.

**Figure 2.**
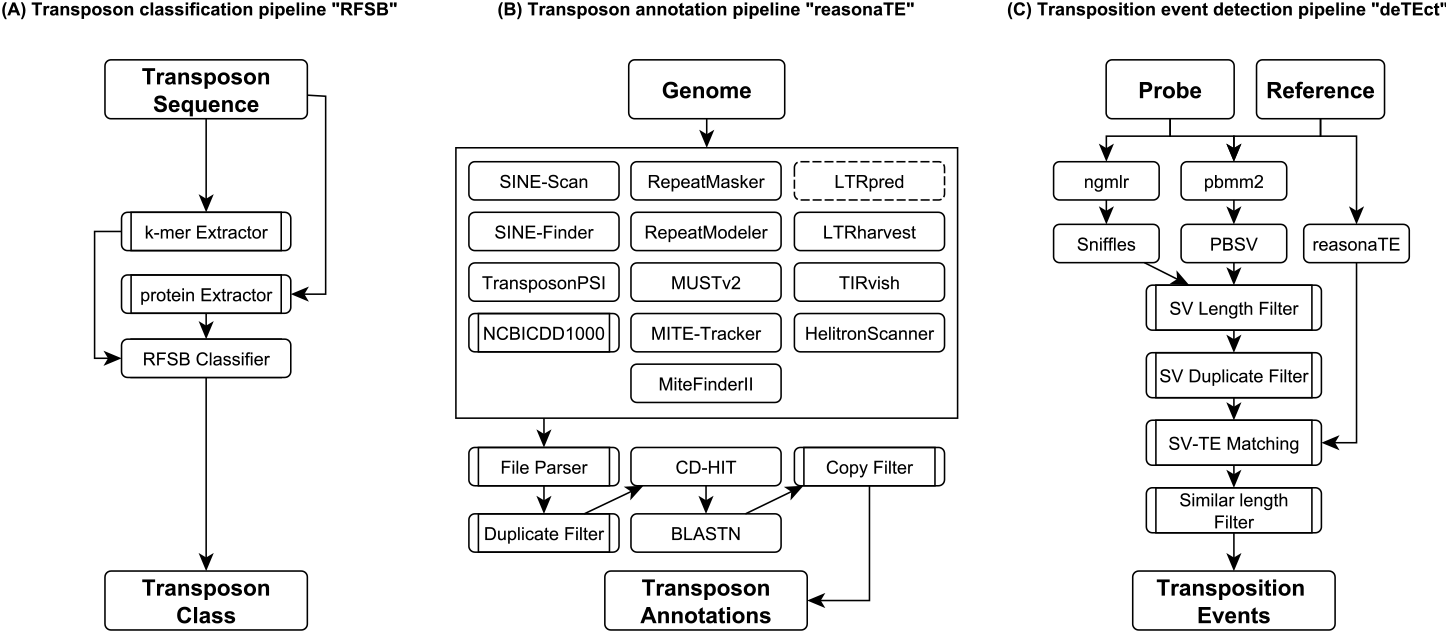
Three pipelines of the TransposonUltimate framework. (A) Given the nucleotide sequence of a transposon, relative k-mer frequencies (for k=2,3,4) and binary protein features are extracted. These features are used by the random forest selective binary classifier (RFSB) to infer the transposon’s class. (B) Published transposon and protein annotation tools are applied to a given genome. Resulting annotations are filtered, merged and clustered using CD-HIT. Then, BLASTN is used to find additional full-length copies. (C) Sequencing reads obtained using a long-read technology from a probe genome are aligned onto a reference genome using ngmlr and pbmm2. Next, the alignments are used to discover structural variants. After filtering the structural variants, they are matched to the transposon annotations to detect transposition events.

## 2 Materials and methods

### 2.1 Transposon classification module, RFSB

Given a nucleotide sequence that is considered to be a transposon, the goal is to determine the class of a transposon according to a given taxonomy. This task is a hierarchical classification problem, meaning the classifier needs to identify multiple classes that stand in a relationship described by a taxonomic hierarchy. The design of the classification module includes several aspects; choosing a transposon database, feature selection, model structure, training strategy, model implementation, evaluation and benchmarking.

The classifiers considered here are supervised learning algorithms, and consequently their performance is limited by the data used for training. Previous studies used small transposon sequence databases, each with different taxonomic schemes, which does not allow for a direct comparison. Therefore, we created *TransposonDB* (Fig. 3, File **F1**), a large collection of transposon sequences that consists of ten databases: ConTEdb [53] (http://genedenovoweb.ticp.net:81/conTEdb/index.php), DPTEdb [54] (http://genedenovoweb.ticp.net:81/DPTEdb/browse.php?species=cpa&name=Carica_papaya_L.), mipsREdat-PGSB [55] (https://pgsb.helmholtz-muenchen.de/plant/recat/index.jsp), MnTEdb [56] (http://genedenovoweb.ticp.net:81/MnTEdb1/), PMITEdb [57] (http://pmite.hzau.edu.cn/download_mite/), RepBase [58] (https://www.girinst.org/repbase/) ^[1]^, RiTE [59] (https://www.genome.arizona.edu/cgi-bin/rite/index.cgi), Soyetedb [60] (https://www.soybase.org/soytedb/#bulk), SPTEDdb [61] (http://genedenovoweb.ticp.net:81/SPTEdb/browse.php?species=ptr&name=Populus_trichocarpa) and TrepDB [62] (http://botserv2.uzh.ch/kelldata/trep-db/downloadFiles.html). To create the database, the taxonomies were unified, duplicates were dropped and several filter rules were applied (Table S1). Filtering included the removal of sequences with no label, the exclusion of fragments, contigs, satellites and RNA sequences. Moreover, only sequences with a length greater than 100bp and those including at least once each of the letters ‘A’,’C’,’G’ and ‘T’ were kept. To the best of our knowledge, this is the largest database of transposon sequences available. Since TransposonDB covers all relevant Eukaryotic kingdoms, it allows for the training and evaluation of a robust, cross-species hierarchical classification model (Table S2 + S3). Moreover, the database is balanced and covers sufficient examples for all taxonomic nodes (Table S4).

**Figure 3.**
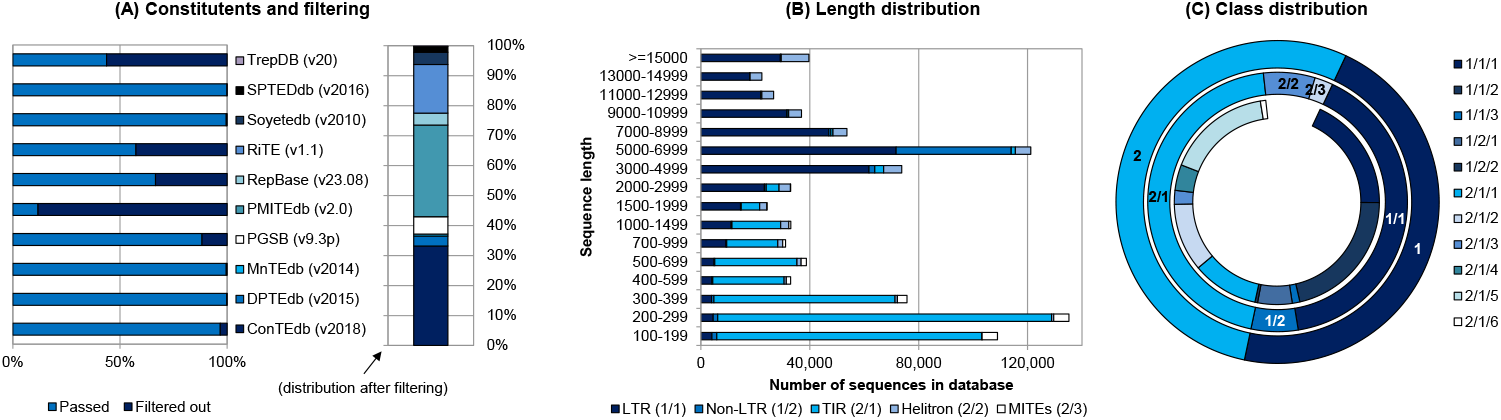
Summary statistics for the TransposonDB. (A) Ten publicly available transposon databases were filtered and combined. Sequences with no (valid) class label, fragments, contigs, satellites, RNA, shorter than 100 bp were filtered out. Moreover, duplicates were dropped when merging. Taxonomic schemes by different databases were unified. (B) The length distribution of sequences in the databases reveals that most DNA transposons are shorter than 500 bp, while most retrotransposons are longer than 3,000 bp. However, Helitrons are significantly longer than other DNA transposons. (C) TransposonDB is balanced in terms of class occurrence, although ERV (1/1/3), SINE (1/2/2) and Novosib (2/1/6) transposons occur rarely.

We selected the combination of relative k-mer frequencies and binary protein features for our classifier. Relative k-mer frequencies represent the number of occurrences of a k-mer within a sequence divided by the number of times it would appear if the sequence consisted of this k-mer only. Protein features are binary, indicating the presence of a certain protein domain in the sequence. The feature vector consists of k-mer frequencies (*k* = 2, 3, 4) and 169 selected domains from NCBI CDD [63] covering class-specific transposons (Table S5). RPSTBLASTN (v2.10.1) was used to annotate the conserved domain models at an e-value of 5.0 as it performed best in terms of classification performance (Fig. S1 A-B). In addition, two model structures were explored. The binary structure employs binary classifiers for each node (= transposon class) of the taxonomy. After inference of the binary classifiers, the taxonomic class can be determined by choosing the most probable node at each stage. The multilabel structure employs a multilabel classifier for each parent node of the taxonomy with *n*+1 classes representing the taxonomic child classes and −1 (return scenario). After inference, the taxonomic class can be determined by choosing the most probable child node at each stage or to return to a higher level and then choose the second most probable child node at that stage. Moreover, we explored two training strategies. The comprehensive training strategy trains each classification node with the whole training set, while the selective training strategy trains each classification child node with a training set that was activated by the parent node. All training strategies, model structures and feature generation were implemented in Python (v3.6.9). Models implementing random forests, AdaBoost, logistic regression, SVM and Naive Bayes from the machine learning package scikit-learn (v0.23) [64] were explored. Random forest consistently yields the highest classification performance (Fig. S2). Based on these results, we propose a **r**andom **f**orest classifier with a **s**elective training strategy on a **b**inary model structure, *RFSB*.

Previous transposon classification studies use different performance measures, taxonomies, training and testing sets, making it hard to compare them. To evaluate the performance, we consider three perspectives. The first perspective is based on hierarchical precision and recall, meaning it considers the whole taxonomy, as proposed in [65]. The second perspective evaluates for different taxonomic levels and the third perspective captures the classification performance of single classes. We benchmark RFSB againts TERL [29], TopDown [24], NLLCPN [27], HC LGA [33] and HC GA [31], as their published code allowed for reproduction. To ensure a fair comparison, source codes were partially modified to allow the training and evaluation of these models on the taxonomy used in our work and TransposonDB.

### 2.2 Transposon annotation module, reasonaTE

Given an assembled genome, the goal of the annotation module is to find all transposon occurrences and their locations. Our *reasonaTE* pipeline produces rich annotations, including transposon mask regions (union of all annotated base pairs) as well as transposon annotations, classification, structural and protein features. This is achieved by combining the advantages of thirteen published transposon annotation tools covering different annotation approaches and transposon classes: RepeatMasker (http://www.repeatmasker.org/), RepeatModeler (http://www.repeatmasker.org/RepeatModeler/), LTRharvest [66] (https://www.zbh.uni-hamburg.de/forschung/gi/software/ltrharvest.html) and TIRvish [67] (http://genometools.org/tools/gt_tirvish.html) are available as Conda packages. Moreover, we created Conda packages for SINE-Finder [68] (http://www.plantcell.org/content/suppl/2011/08/29/tpc.111.088682.DC1/Supplemental_Data_Set_1-sine_finder.txt), SINE-Scan [69] (https://github.com/maohlzj/SINE_Scan), HelitronScanner [42] (https://sourceforge.net/projects/helitronscanner/files/), MUSTv2 [70] (http://www.healthinformaticslab.org/supp/resources.php), MiteFinderII [71] (https://github.com/jhu99/miteFinder) and MITE-Tracker [72] (https://github.com/INTABiotechMJ/MITE-Tracker) to make them accessible and to facilitate their installation. Also, we include the output files of LTRpred [73] (https://hajkd.github.io/LTRpred/articles/Introduction.html) into the pipeline, as this tool provides high quality annotations, but is available as a Docker image only. As the tools have different output formats, we developed a parser module to convert all outputs to GFF3 format.

After running the annotation tools, additional copies of the identified transposons are searched using the clustering tool CD-HIT (v4.8.1) [74, 75] and BLASTN (v2.10.1). For the annotation of transposon-characteristic proteins, we have created a Conda packaged version of TransposonPSI (http://transposonpsi.sourceforge.net/), and we also use the protein domains from NCBI CDD for this task. Using TransposonDB, NCBI CDD and RPSTBLASTN, we selected the 1,000 most frequently occurring protein domains that are characteristic to transposons (File **F2**). As an application, here we annotate the genome *MSU7* of *Oryza sativa subspecies japonica* (http://rice.plantbiology.msu.edu/index.shtml), the genome *DAOM197198* of *Rhizophagus irregularis* (https://www.ncbi.nlm.nih.gov/bioproject/?term=PRJDB4945) [76], three reference genomes *VC2010* (https://www.ncbi.nlm.nih.gov/bioproject/?term=PRJEB28388), *N2* (https://www.ncbi.nlm.nih.gov/bioproject/?term=PRJNA13758), *CB4856* (https://www.ncbi.nlm.nih.gov/bioproject/?term=PRJNA275000) and twenty novel wild type strains [77] of *Caenorhabiditis elegans* (Table S6).

### 2.3 Transposition event detection module, deTEct

Given an assembled reference genome and sequenced probe genome reads, the goal is to identify transposition events that are manifested as structural variants. This requires both a list of SVs and annotation of TEs as inputs. We employ the structural variant caller Sniffles on ngmlr [78] alignments and PBSV (https://github.com/PacificBiosciences/pbsv) structural variant caller on pbmm2 alignments (https://github.com/PacificBiosciences/pbmm2). Moreover, the TE annotations are generated using the proposed reasonaTE pipeline mentioned before.

SVs are filtered twice. First, variants shorter than 50 bp or longer than 1% of the genome length were excluded. Second, duplicate structural variants of the same type are merged. Consecutively, the remaining variants and TE annotations are matched and finally reported if their length corresponds to each other. Transposon annotations were matched to structural variants if they intersected for at least 10% and their length was similar by a threshold of 50%. We chose to do so, as structural variant callers and transposon annotators have an uncertainty regarding exact locations. We therefore consider a similar length more important than a high overlap. The proposed deTEct pipeline is applicable to long-read sequencing technologies, and it has been tested with both PacBio and OxfordNanopore data.

## 3 Results

### 3.1 RFSB outperforms other transposon classifiers

We benchmarked our RFSB method against other transposon classifiers, and the results show that it has the highest sensitivity and specificity (Fig. 4 A, Table S7). TE Learner [20] has the lowest reported performance, while the other methods have similar F1 scores. However, this comparison is based on reported numbers from different studies with different evaluation schemes, taxonomies and datasets for training and testing. For a more fair comparison some of the tools were applied to the subset of TranspsonDB which includes RepBase and PGSB (Fig. 4 B). The comparison of the results reveals large discrepancies. Surprisingly, TERL and TopDown have a performance which is worse than random guessing, and closer inspection of the outputs from NLLCPN reveals that it has learned a constant distribution rather than a relationship between sequences and classes.

**Figure 4.**
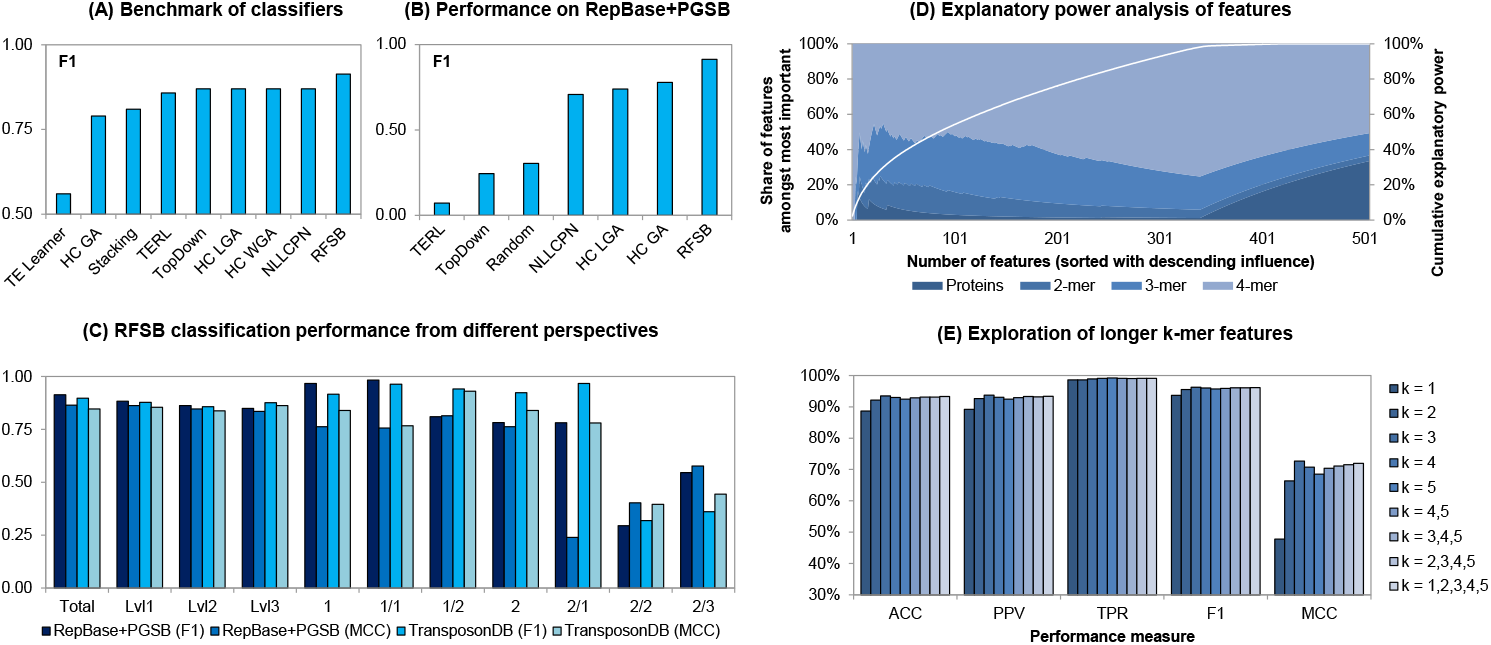
Evaluation of the RFSB classifier. (A) Benchmark of different transposon classifiers by reported numbers in publications. (B) The performance of selected, reproducable classifiers applied on RepBase+PGSB database using the taxonomy in Fig. 1 A. Reported numbers represent performances from a total perspective. (C) RFSB classification performances from total, taxonomic level and class perspective. (D) Analysis of each feature’s contribution to classifier’s explanatory power. The white line shows the cumulative explanatory power. (E) Analysis of different k-mer features in combination with protein features for a binary classifier differentiating between class 1 and 2 transposons. All values presented were calculated as average across a tenfold cross validation.

A detailed analysis of the classification performance of RFSB across different taxonomic levels and classes reveals a small decrease in performance when considering deeper taxonomic levels (Fig. 4 C). Underrepresented classes, e.g. Helitrons and MITEs, perform worse, and the results are consistent for both F1 and MCC scores. Moreover, for some classes the performance of RFSB on the large, cross-species TransposonDB is better than for the more homogeneous subset of RepBase and PGSB, which suggests that it is robust, generalisable, and applicable to different species. An inspection of the most informative features (File **F3**) shows that longer k-mer features contribute the most to the classification performance, while protein domains have a smaller share amongst the most contributing features (Fig. 4 D). This motivated the exploration of longer k-mer features, but we did not find any significant increase of the performance when using 5-mers (Fig. 4 E).

### 3.2 The ensemble strategy reasonaTE finds more transposons and reduces bias

Next, we evaluated the ability of our reasonaTE pipeline to identify TEs in the genomes of three different species (Fig. 5 A-B, File **F4**). The TE content of almost 21% for *Caenorhabditis elegans* is higher than previously reported values of 12% [8], 17% [79], and 12-16% [80]. However, as these studies used methods that were biased towards finding specific classes of transposons, it is to be expected that our ensemble strategy finds more TEs. By contrast, the prediction of 33% for *Oryza sativa subs. japonica* is very close to the mean of other reports [81, 82, 83, 84, 85, 86, 87, 88, 89]. The content of 23% in *Rhizophagus irregularis* is close to a previous estimate of 27% [90]. The low variation of transposon content across different strains becomes obvious for the cluster of *Caenorhabditis elegans*. Interestingly, the relative transposon class frequency reveals clear differences across species (Fig. 5 C-D). Similarly, the length distributions (Fig. 5 E-G) exhibit substantial differences between transposons of the same class found in different species. Helitrons in particular vary in length as was observed before [91].

**Figure 5.**
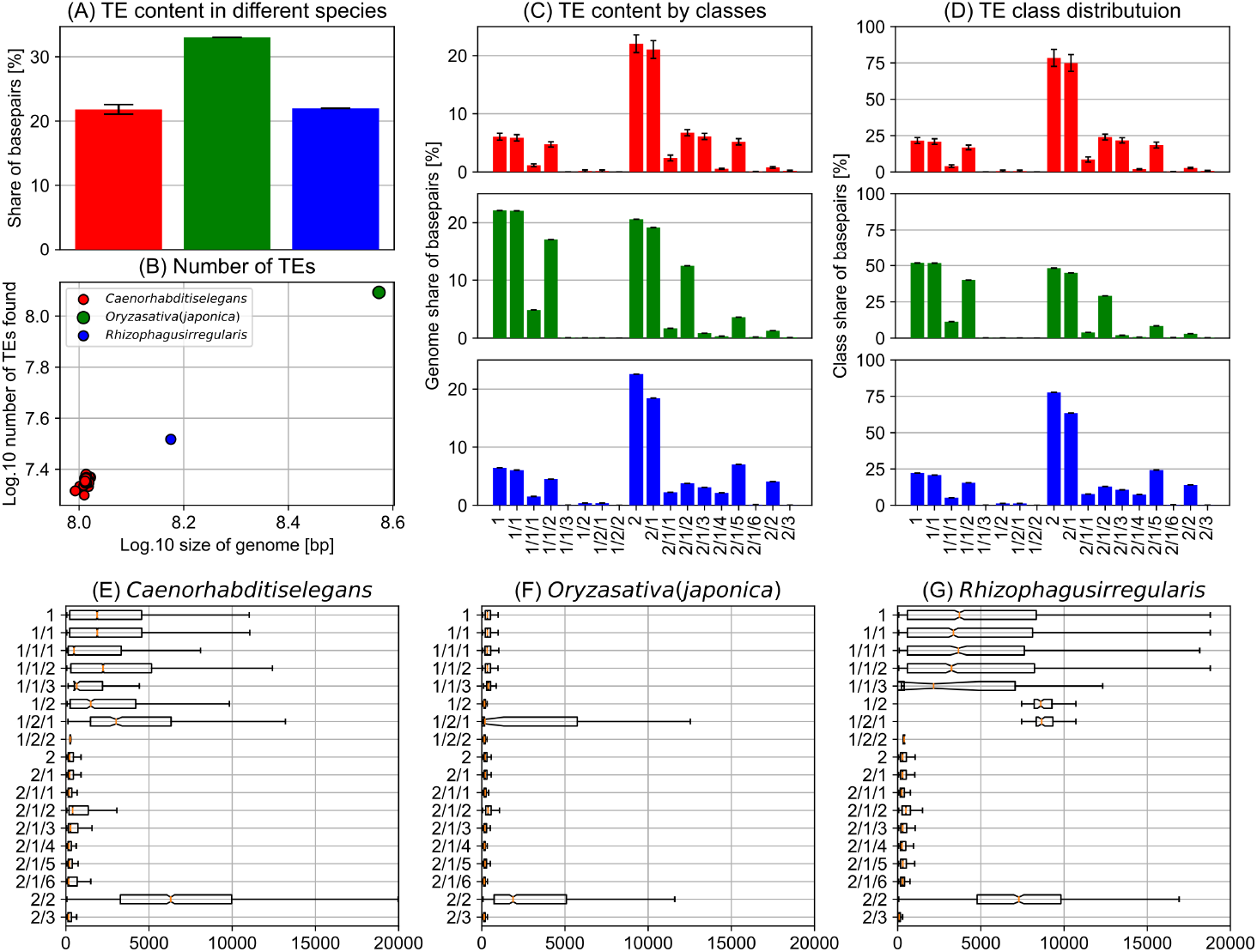
reasonaTE results for three species. The colors used in this figure represent *Caenorhabditis elegans* (red), *Oryza sativa subs. japonica* (green) and *Rhizophagus irregularis* (blue). (A) The average TE content of different species. The TE content is calculated as ratio of the sum of all basepairs part of the transposon region mask and the total genome size. The whiskers represent standard deviations. (B) The dot size represents the TE content as reported in the first panel, and the figure shows a linear relationship between genome size and the total number of transposons found. (C) Average TE content by transposon classes. The values were calculated by dividing the sum of the lengths of all transposons of a specific class by the total genome length. The whiskers represent the standard deviation. (D) The class distribution across all TEs based on the number of elements. (E-G) The transposon length distribution by classes for the three species. The boxes cover 25% to 75% percentiles, including the orange bar at the 50% percentile. The length of whiskers amounts to 150% of the interquartile range.

In concordance with [92] and [93], the share of Helitrons amounts to almost 2% of the *Caenorhabditis elegans* genome. Moreover, the majority of the transposons are TIR DNA transposons, as reported by [94, 95, 79]. Contrary to previous studies [80, 96, 97], we mainly find hAT, CMC and Novosib transposons to be present in the *Caenorhabditis elegans* genome rather than Tc1-Mariner transposons. Our findings for the rice genome are consistent with previous findings. The high frequency of Gypsy (class 1/1/2) compared to other LTR (class 1/1) and non-LTR (class 1/2) was reported in *Oryza sativa subs. japonica* [87]. Moreover, the small share of MITEs, up to 2%, is similar to the previously reported share of 4% [89]. A previous study [44] found that class 1 transposons have a larger share (25%) than class 2 transposons (20%) and the frequencies for the subclass level (LTR 23.5% and non-LTR 2%, TIR 17.5% and Helitrons 3.6%) match our findings. Inspection of the annotation density across the chromosomes revealed a characteristic concentration at the arms for *Caenorhabiditis elegans* (Figure S3), consistent with the higher densities observed for other variants [79, 98, 99, 100, 101, 80].

The comparison of different annotation tools reveals that reasonaTE provides more unbiased results (Figure S4) as none of the other methods find more than 31.8% of the TEs reported by reasonaTE. In addition, the analysis shows that around 40% of the repetitive elements found by RepeatMasker and RepeatModeler were confirmed as transposons using our approach. Moreover, the transposon characteristic protein annotations by TransposonPSI and the 1,000 most frequently occurring proteins from NCBI CDD intersect significantly with reasonaTE’s transposon annotations. The analysis also reveals large overlap between some tools, e.g. MUSTv2 & MITE-Tracker, LTRpred & LTRharvest, and SINE-Finder with all other tools.

Closer inspection of the class composition of the TEs found for *Caenorhabditis elegans* confirms the advantages of the ensemble technique of reasonaTE (Figure S5). None of the tools is able to find the same share of TEs on its own as the ensemble. Moreover, we find that tools that were designed to identify a specific transposon class annotate TEs from different classes as well.

### 3.3 29,554 transposition event candidates were observed analyzing 20 wild type strains of *Caenorhabditis elegans* using deTEct

Finally, we applied the deTEct pipeline to 20 whole genome assemblies of wild type strains of the nematode *Caenorhabditis elegans*. Each strain was compared to the two reference genomes *VC2010* and *CB4856* (Fig. 6 A, Table S8, File **F5**). As expected, the newly sequenced genomes of these two strains have almost no transposition events when compared to their reference. Closer inspection of the transposon and transposition event densities reveals that the putative transposition events are primarily located at the ends of the chromosomes (Fig. 6 B) as reported by [79]. From the initial list of SVs, 3.97% were identified as transposition events. However, the list included numerous duplicates or very short variants that were subsequently filtered out. Consequently, we find that after filtering, 7.37% of all SVs are caused by transposition events.

**Figure 6.**
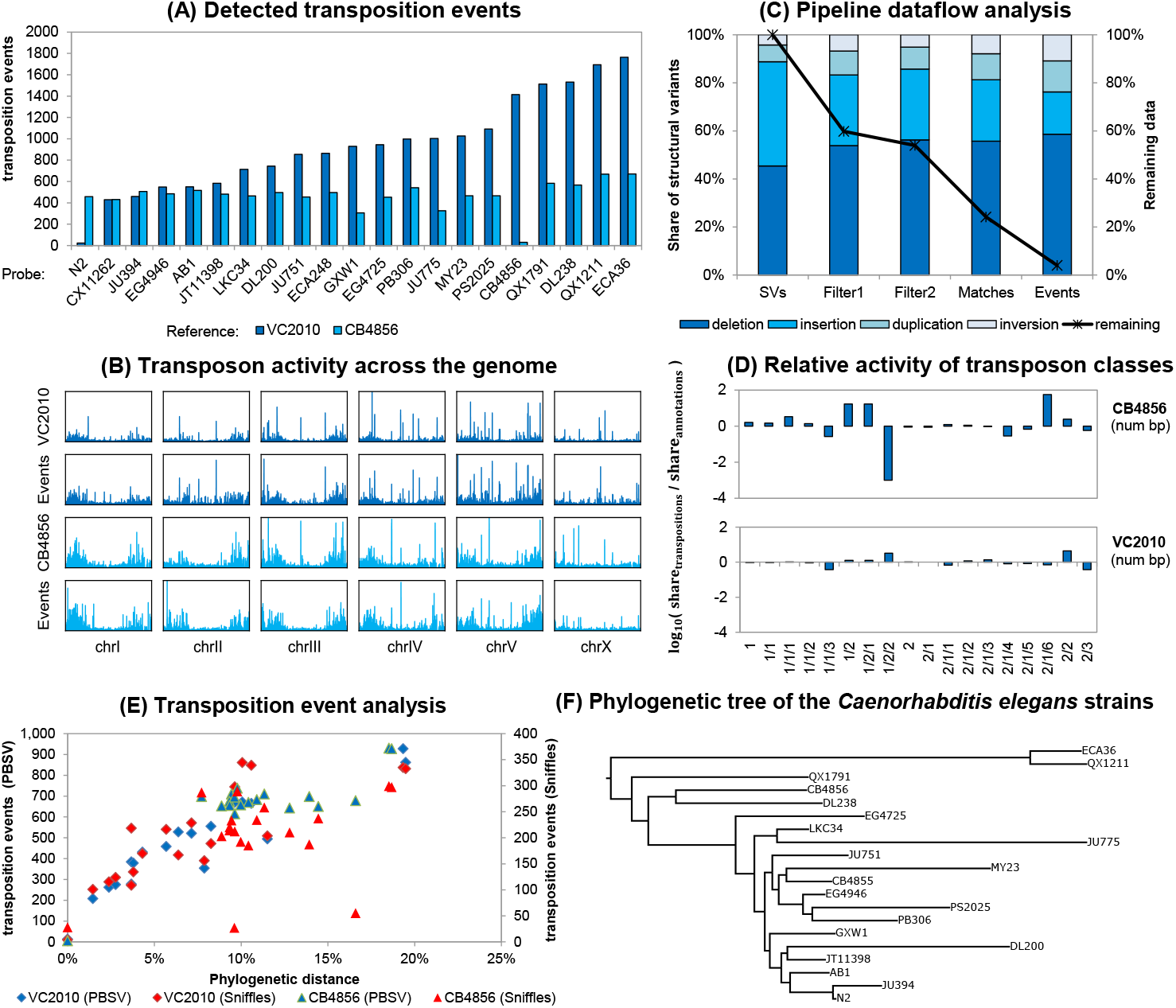
deTEct results and discovered transposition events. (A) Results show the number of detected transposition event candidates by probe strain for both reference genomes *VC2010* and *CB4856*. (B) The transposon activity in the *Caenorhabditis elegans* genome by chromosomes. The first row shows the density of transposon annotations in *VC2010*. The second row shows the density of transposition events. The following two rows represent results for *CB4856*. For all autosomal chromosomes we identify a characteristic pattern of transposon activity at the ends of chromosomes. (C) Dataflow analysis of the pipeline. The diagram shows the share of different structural variant categories at each stage of the pipeline (left y-axis). Deletions make up the largest share of transposition events. Additionally, the share of remaining data is outlined (right y-axis). Approximately 4% of all structural variants initially found are finally identified as transposition events. (D) Helitrons and SINEs are more active relative to *VC2010*, while Novosib are especially active relative to *CB4856*. Relative activity is calculated by the share of a class’ basepairs appearing in transposition events divided by its share of a the classes basepairs in the transposon annotation. (E) A linear relationship between phylogenetic distance and the number of observed transposition events becomes obvious for the *Caenorhabditis elegans* strains for both SV callers PBSV and Sniffles. Phylogenetic distance is calculated as sum of distances in the phylogenetic tree to the last common ancestor. (F) The phylogenetic tree of the *Caenorhabditis elegans* strains. The branch lengths are proportional to the number of polymorphisms that differentiate each pair. Tree based on data from [101].

Most of the transposition events were observed due to deletions (60%) while insertions, duplications and inversions cause the remaining variation (File **F6** + **F7**). One difficulty in interpreting these proportions stems from the known biases of sequencing data [102] which make insertions hard to detect. This results in an elevated number of observations of cut transpositions (deletions), but fewer paste transpositions (insertions). Nonetheless, we find certain classes of transposons to be especially active in the comparisons of probe and reference genomes, such as Helitrons and SINEs relative to *VC2010*, and LINEs and Novosib when compared to *CB4856* (Fig. 6 D, File **F8**). The activity of Helitrons was observed previously [92, 93]. Helitrons were implicated in the divergence of GPCR genes and heat shock elements. Moreover, they are considered to play an important role in evolution [42]. Comparing the two major classes, we conclude that the biggest contribution stems from DNA transposons (82% for VC2010 comparisons and 95% for CB4856 comparisons), similar to the findings in [103].

Moreover, we observe a linear relationship between the number of transposition events found and the phylogenetic distance of the given strains (Fig. 6 E-F, Table S9). The strains *QX1211* and *ECA36* have the largest differences based on transposon data before [80].

## 4 Discussion

Here we present TransposonUltimate, a bundle of three modules for transposon classification, annotation and transposition event detection. Moreover, we present TransposonDB, a database containing more than 891,051 transposon sequences from a wide range of species. Our benchmarks shows that the classification module RFSB outperforms existing methods. Although *RFSB* has a very high accuracy, we believe that performance could be improved by developing species specific classifiers. It would also be helpful to explore new feature representations that strongly correlate to phylogenetic distance metrics.

The annotation module combines existing annotation approaches using an ensemble strategy, and this ensures a less biased outcome than existing methods that tend to favor certain TE classes. The annotation module could be extended by the search for fragmented copies of annotated transposons connected with filters to avoid false positives. Application to three different species revealed that TEs from the same family vary drastically in length. Thus, an important question for future research is to determine to what extent such differences reflect hitherto uncharacterized families, and to what extent the differences correspond to overall sequence divergence.

The detection module enables the identification of transposition events through structural variants in genomes profiled using long-read sequencing technologies. Application of the *deTEct* pipeline to 20 wild type strains of *Caenorhabditis elegans* suggests that transposon events are responsible for 7.37% of structural variants. Although previous studies have argued that transposons are a major driver of structural variation [102], our results suggest that at least for wild isolates of *Caenorhabditis elegans* this is not the case. As additional high quality assemblies become available, it will be interesting to further explore this important question. Moreover, the development of localisation algorithms of target and donor sites of transposons seems a promising add-on for the detection module. Besides, structural variants gathered from whole genome comparison using anchor filtering [104] could be included and compared.

As long-read technologies are becoming more widely used and the number of sequenced genomes rises quickly, there is an urgent need for methods to identify and annotate TEs which correspond to plurality and in some cases a majority of genome sequences. In particular, as more human [105] and other vertebrate (https://vertebrategenomesproject.org/) genomes are profiled using these technologies, TransposonUltimate will be a valuable tool to improve our understanding of the impact of TEs on both traits and diseases.

## 5 Conclusions

Our TransposonUltimate bundle of software tools provides a powerful and user-friendly means of analyzing TEs. In addition to providing highly accurate classifications, our analysis also provides insights as to what features are most informative for predicting TE class. Our ensemble approach to annotation is more unbiased than existing methods that tend to focus on one or a few classes. Finally, our transposition event detection module can take advantage of long-read technologies to identify to what extent TEs underlie SVs.

## Author’s contributions

The study was conceived and designed by KR, CR, EAM and MH. The code was written by KR, and the analyses were carried out by KR and CR. The work was supervised by EAM and MH. KR and MH wrote the manuscript with input from EAM and CR.

## Acknowledgements

We would like to thank Sarah Buddle, Simone Procaccia, Fu Xiang Quah and Alexandra Dallaire for assistance with testing and debugging the software.

## Additional Files

**File F1 : TransposonDB.fasta**

**File F2 : NCBICDD1000 Proteins.txt**

**File F3 : Classification FeatureImportanceAnalysis.csv**

**File F4 : GFF3 files in “PaperSupplements/Annotation/…”**

**File F5 : GFF3 files in “PaperSupplements/Detection/…”**

**File F6 : Detection SVDistribution.csv**

**File F7 : Detection PipelineData.csv**

**File F8 : Detection ClassDistribution.csv**

## Supplements

**Table S1.** Classification: TransposonDB filter rule application. This table shows the number or remaining sequences in the constituents databases of TransposonDB after the application of the different filter rules.

| database | before filtering | #F0<br>no label | #F1.1<br>fragments | #F1.2<br>contigs | #F1.3<br>satellites | #F1.4<br>RNA | #F1<br>all #F1.* | #F2<br>len.100bp | #F3<br>alphabet | #F4<br>all rules |
| --- | --- | --- | --- | --- | --- | --- | --- | --- | --- | --- |
| ConTEdb | 322,705 | 322,702 | 322,702 | 322,702 | 322,702 | 322,702 | 322,702 | 317,109 | 317,995 | 312,402 |
| DPTedB | 31,340 | 31,326 | 31,326 | 31,326 | 31,326 | 31,326 | 31,326 | 31,285 | 31,325 | 31,284 |
| PGSB | 61,730 | 61,729 | 61,729 | 61,729 | 61,529 | 60,813 | 55,616 | 60,757 | 61,397 | 54,488 |
| MnTEdb | 5,925 | 5,912 | 5,912 | 5,912 | 5,912 | 5,912 | 5,912 | 5,902 | 5,912 | 5,902 |
| PMITEdb | 2,449,127 | 357,134 | 313,211 | 313,211 | 313,211 | 313,211 | 313,211 | 298,777 | 302,529 | 288,104 |
| RepBase | 56,403 | 53,880 | 51,674 | 51,674 | 51,674 | 51,648 | 51,674 | 51,455 | 37,796 | 37,563 |
| RiTE | 265,549 | 264,022 | 242,216 | 156,195 | 241,847 | 240,509 | 154,232 | 152,746 | 154,058 | 152,575 |
| Soyetedb | 38,664 | 38,603 | 38,603 | 38,603 | 38,603 | 38,603 | 38,603 | 38,519 | 38,601 | 38,517 |
| SPTEDdb | 18,413 | 18,408 | 18,408 | 18,408 | 18,408 | 18,408 | 18,408 | 18,402 | 18,408 | 18,402 |
| TrepDB | 4,162 | 3,910 | 1,874 | 3,910 | 3,910 | 3,910 | 1,870 | 3,669 | 3,869 | 1,822 |
| TransposonDB |  |  |  |  |  |  |  |  |  | 891,051 |

**Table S2.** Classification: TransposonDB sequences across biological domains. This table shows the number of sequences in TransposonDB by the source database and biological domains.

| databases | ConTEdb | DPTEdb | mipsREdat-PGSB | MinTEdb | PMTEdb | RepBase23.08 | RiTE | Soyetedb | SPTEDdb | TrepDB |
| --- | --- | --- | --- | --- | --- | --- | --- | --- | --- | --- |
| Eukaryota | 312,334 | 29,452 | 52,691 | 3,154 | 282,500 | 36,681 | 129,911 | 36,429 | 6,733 | 1,068 |
| Animalia | 0 | 0 | 0 | 0 | 0 | 21,854 | 0 | 0 | 0 | 20 |
| Chromista | 0 | 0 | 0 | 0 | 256 | 988 | 0 | 0 | 0 | 74 |
| Fungi | 0 | 0 | 0 | 0 | 0 | 1,961 | 0 | 0 | 0 | 354 |
| Plantae | 312,334 | 29,452 | 52,691 | 3,154 | 282,244 | 11,721 | 129,911 | 36,429 | 6,733 | 620 |
| Protozoa | 0 | 0 | 0 | 0 | 0 | 150 | 0 | 0 | 0 | 0 |
| UnicellularFlagellate | 0 | 0 | 0 | 0 | 0 | 7 | 0 | 0 | 0 | 0 |
| Prokaryota | 0 | 0 | 0 | 0 | 0 | 54 | 0 | 0 | 0 | 6 |
| Virus | 0 | 0 | 0 | 0 | 0 | 31 | 0 | 0 | 0 | 0 |
| Total | 312,334 | 29,452 | 52,691 | 3,154 | 282,500 | 36,766 | 129,911 | 36,429 | 6,733 | 1,074 |

**Figure S1.**
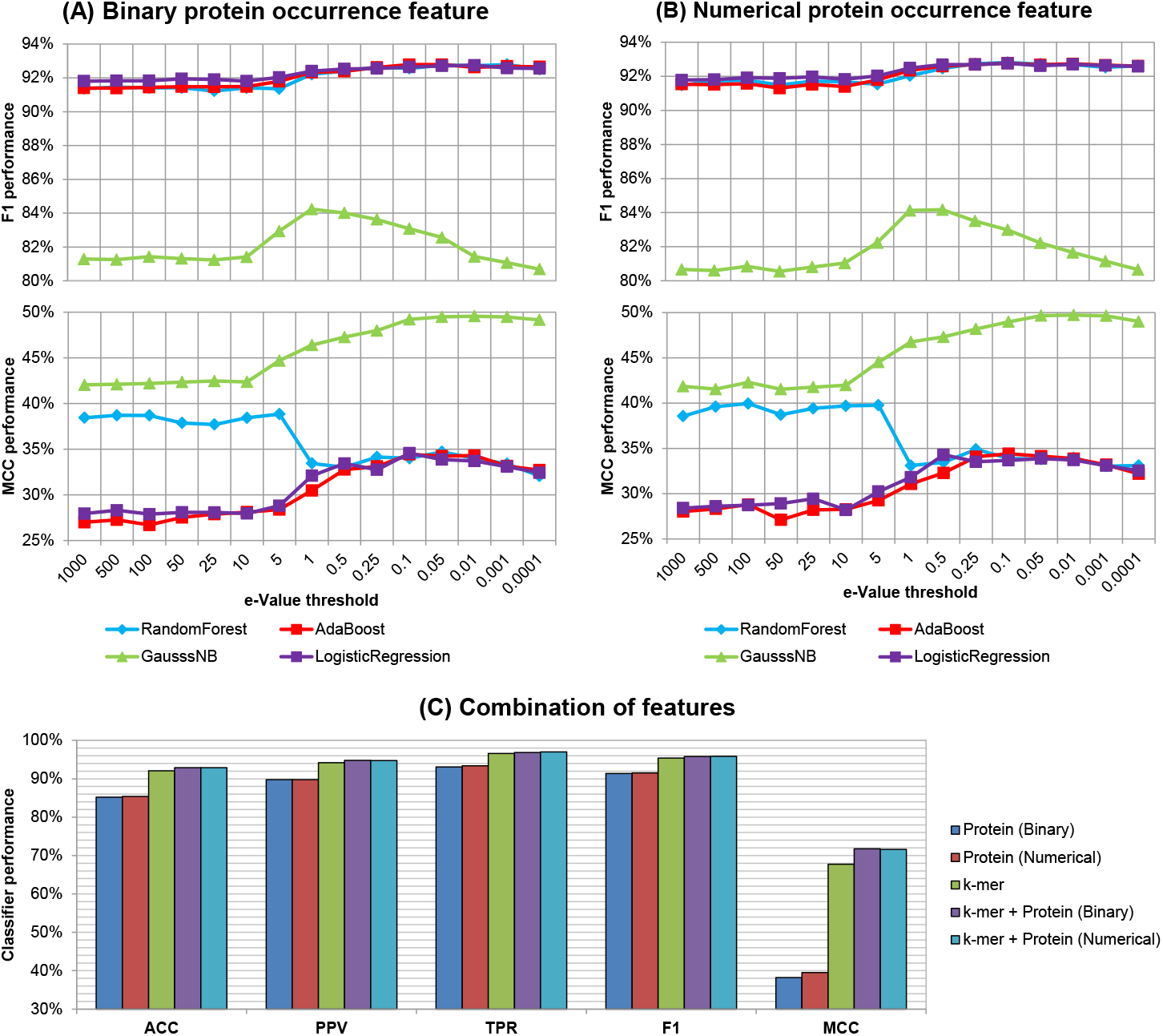
Classification: feature design experiments. All performance measures reported as average across a tenfold cross validation (in %) on RepBase+PGSB dataset for a classifier distinguishing class I and II transposons. (A) Performance of standard classifiers using the binary protein feature for different e-thresholds. This feature is either zero or one depending on whether the protein domain is detected for given e-threshold by BLASTN. (B) Performance of standard classifiers using the numerical protein features for different e-thresholds. This feature represents the number of times the protein domain is detected for given e-threshold by BLASTN. (C) Combinations of the protein features with relative k-mer frequency for a random forest classifier.

**Figure S2.**
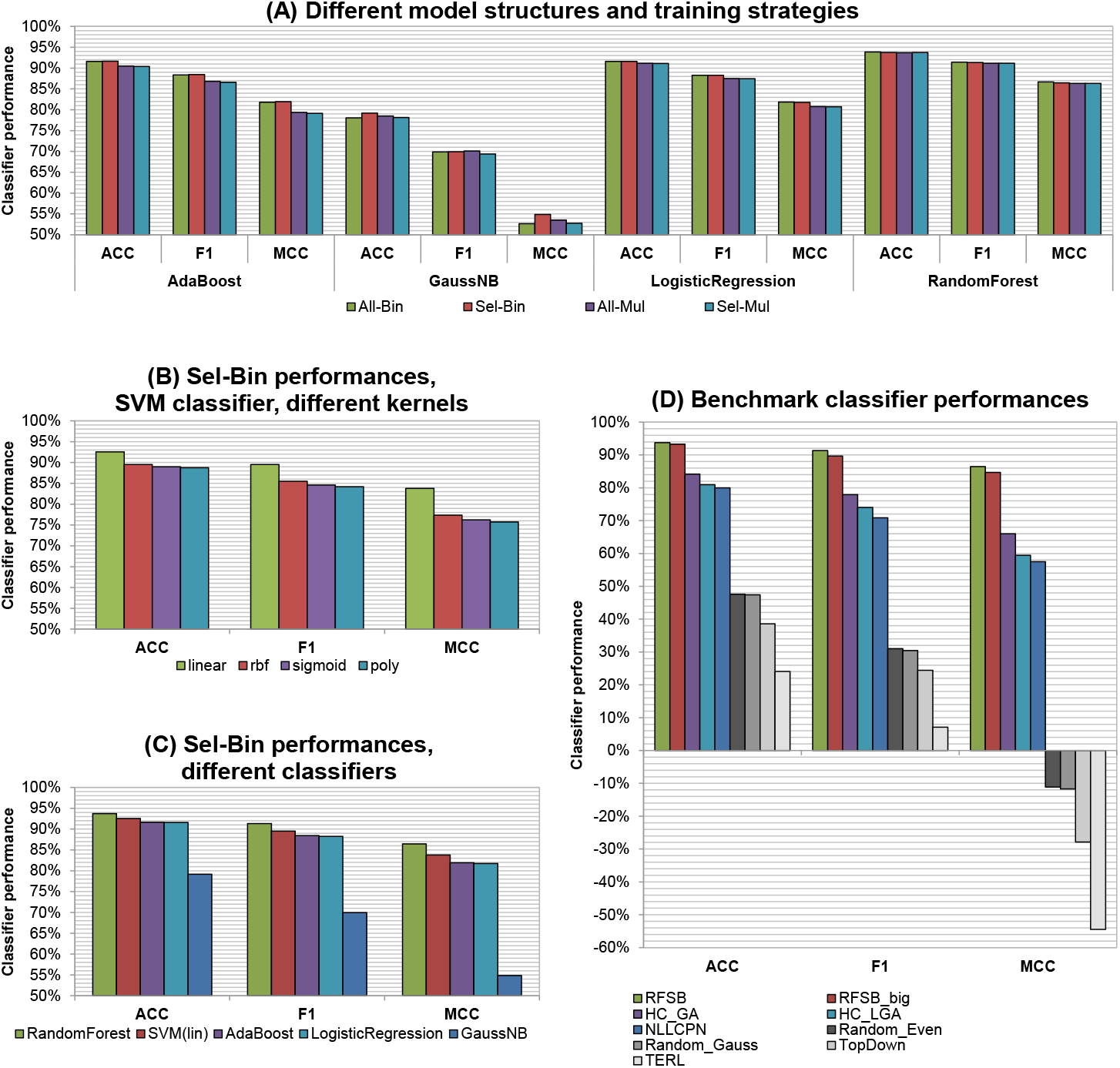
Classification: structure, strategy and model design experiments. All performance measures reported as average across a tenfold cross validation (in %) on RepBase+PGSB from an overall perspective. Panel (A) exhibits the performance of standard classifiers for different model structure and training strategy combinations. Panel (B) shows the performance of SVM classifier model for different kernel functions. Panel (C) summarises the performances of standard classifiers and best SVM classifier for the selective binary (Sel-Bin) combination. Panel (D) compares the proposed “RFSB” (Random Forest Selective Binary) approach with existing benchmark classifiers. In addition, RFSB trained on TransposonDB is reported as well.

**Figure S3.**
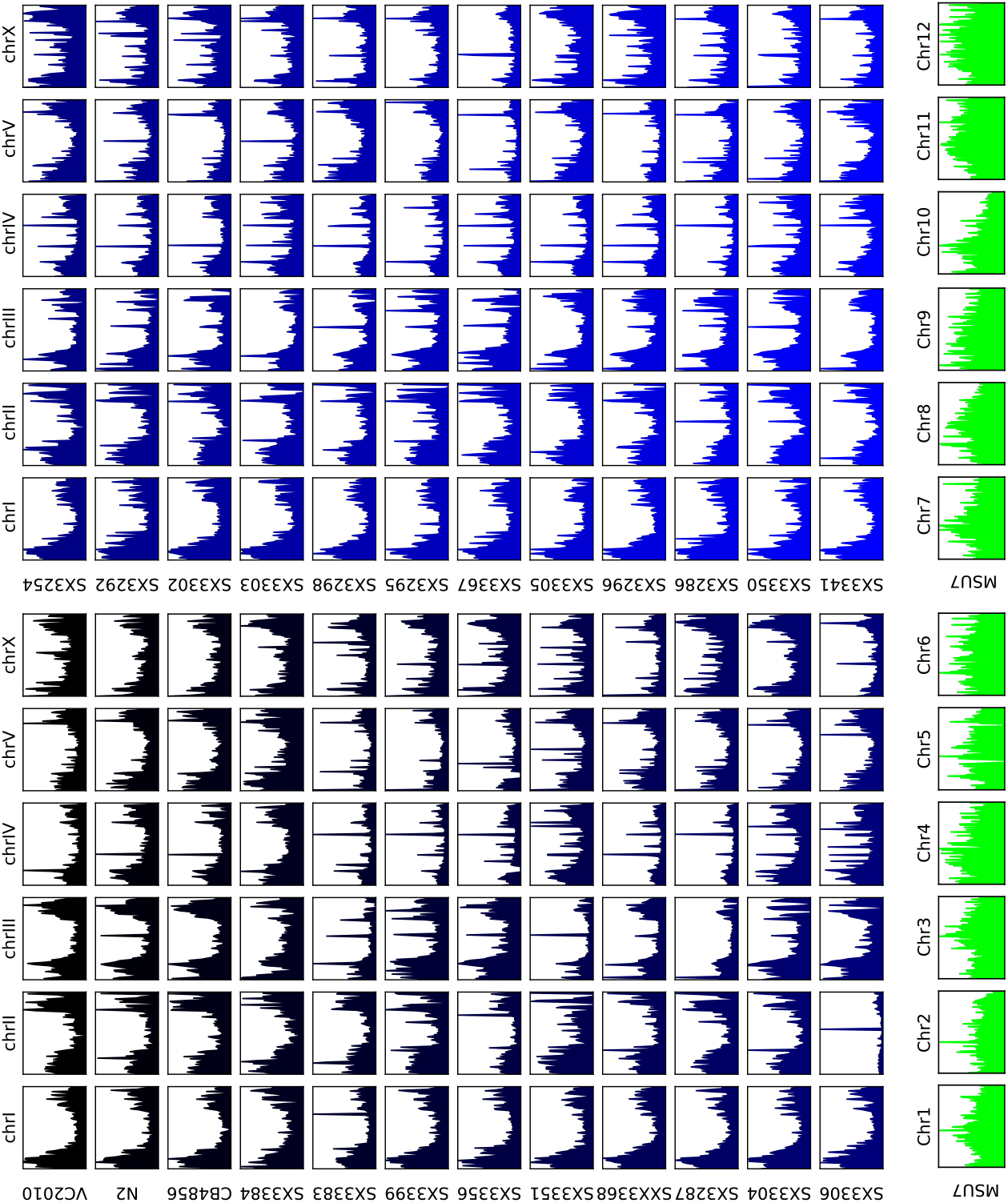
Annotation: transposon annotation density plots. This diagram shows the density of transposon annotations across the chromosomes of the different *Caenorhabditis elegans* and the *Oryza sativa subs. japonica* genomes.

**Figure S4.**
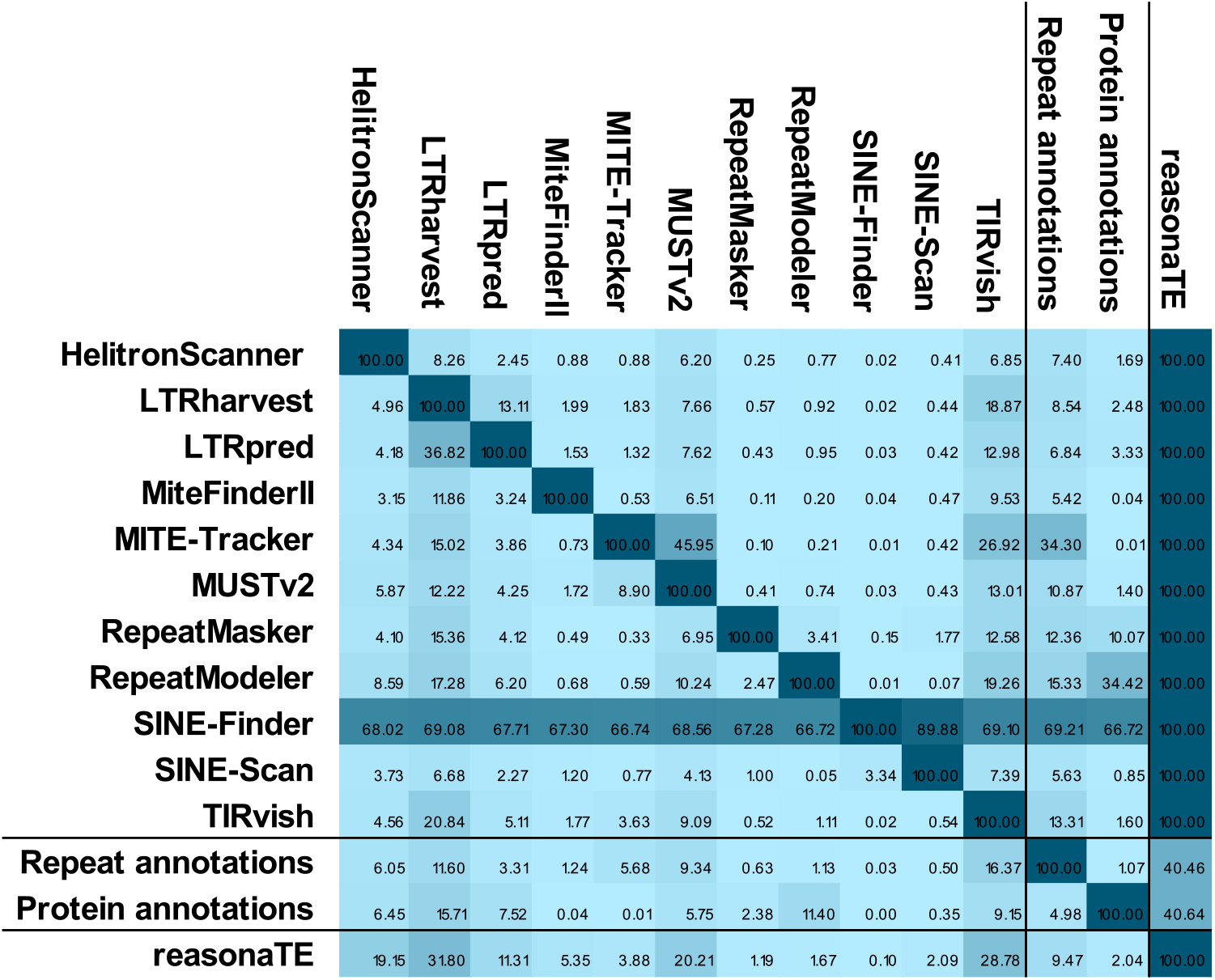
Annotation: tool intersection heat map. This heat map presents the intersection of annotations from different tools. The content of cell in row i and column j represents the number of intersecting basepairs of annotations from tools i and j, divided by the total number of basepairs in annotations of tool row i. Numbers are reported in percent, as average across the three reference genomes *VC2010*, *N2* and *CB4856*. Repeat annotations are the combination of annotated repeats by RepeatMasker and RepeatModeler. Protein annotations are the combination of annotated transposon characteristic proteins by NCBICDD1000 and TransposonPSI. The colour of a cell represents its value. Darker colours represent values closer to 100%, while lighter colours represent values closer to 0%.

**Figure S5.**
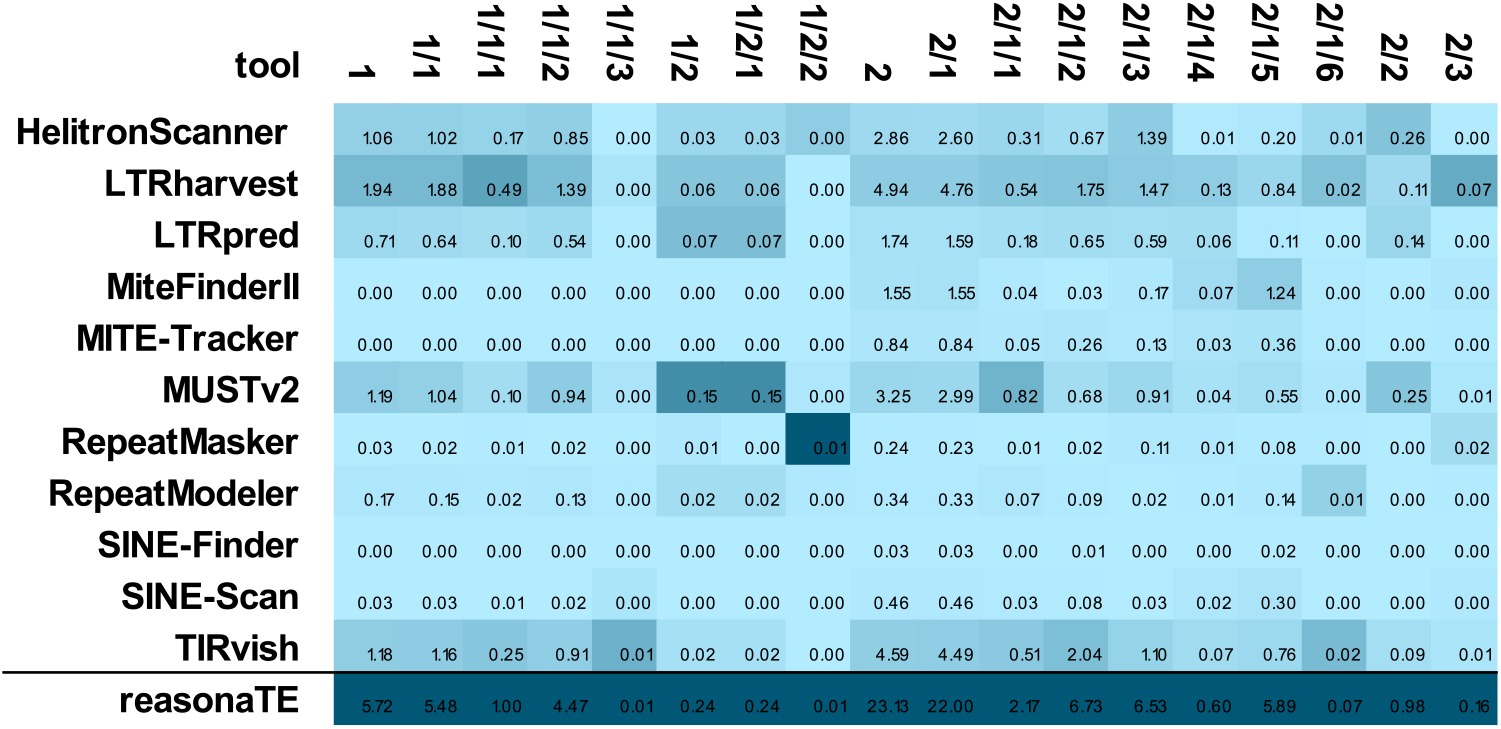
Annotation: tool class heat map. This heat map presents the share of annotated transposons by different transposon classes. The content of a cell represents the number of basepairs of a specific tool’s annotations related to a specific class divided by the total genomes length. Numbers are reported in percent, as average across the three reference genomes *VC2010*, *N2* and *CB4856*. The colour of a cell represents its value. The darker the colours, the more a tool was able to capture most of the transposons that the ensemble (reasonaTE) found for this specific class.

**Table S3.**
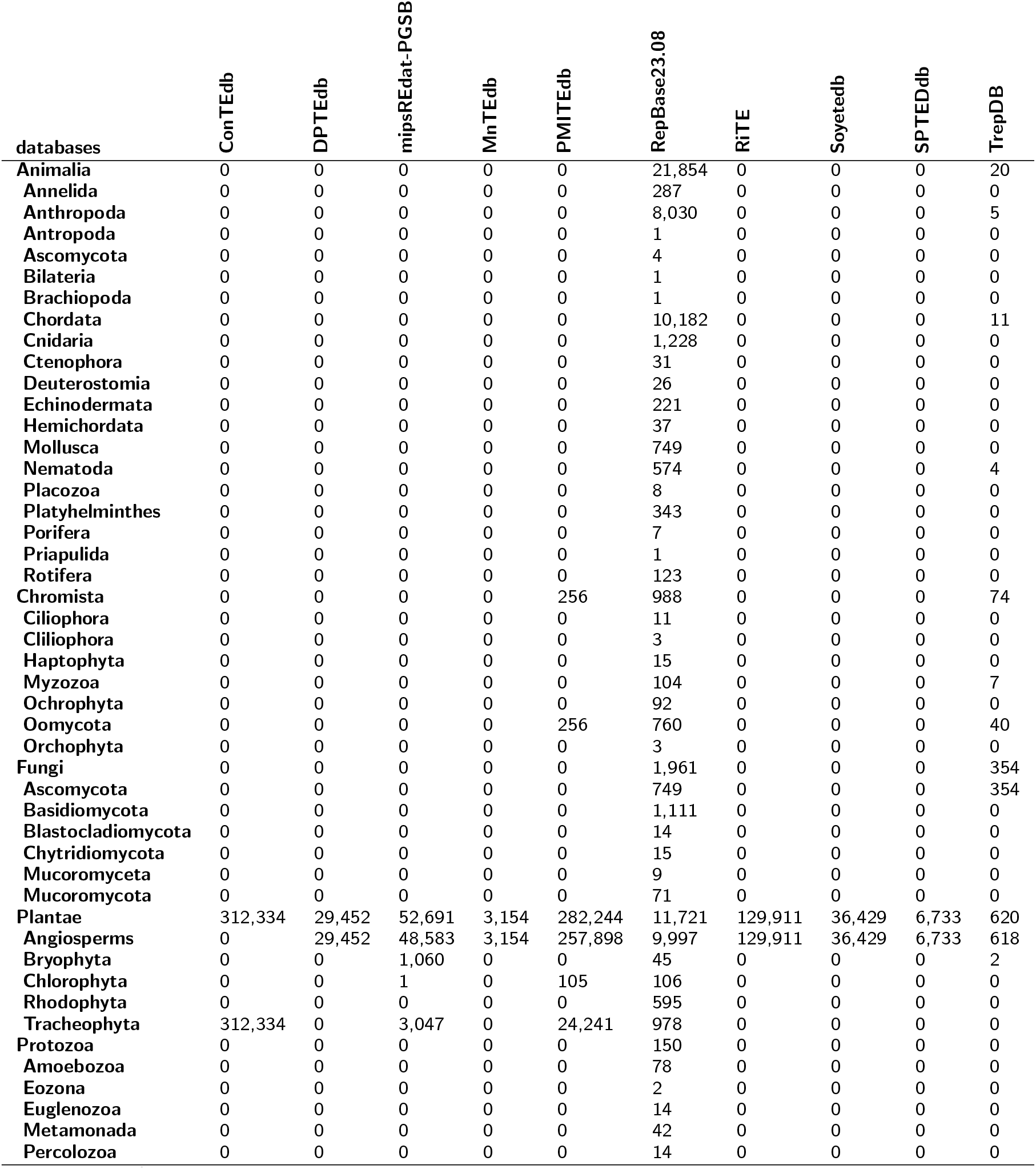
Classification: TransposonDB sequences across eukaryotic kingdoms. This table shows the number of sequences in TransposonDB by the source database and biological kingdoms.

**Table S4.** Classification: TransposonDB sequences across transposon classes. This table shows the number of sequences in TransposonDB by the source database and transposon classes.

| database | TransposonDB | ConTEdb | DPTeddb | mipsREdat-PGSB | MnTEdb | PMITEdb | RepBase23.08 | RIITE | Soyetedb | SPTEdDb | TrepDB |
| --- | --- | --- | --- | --- | --- | --- | --- | --- | --- | --- | --- |
| 1 | 412,975 | 252,236 | 23,832 | 49,853 | 1,280 | 0 | 25,908 | 23,508 | 30,538 | 4,972 | 848 |
| 1.1 | 360,618 | 210,831 | 22,343 | 48,186 | 1,267 | 0 | 21,542 | 20,670 | 30,404 | 4,769 | 606 |
| 1.1.1 | 122,715 | 81,329 | 5,010 | 11,059 | 600 | 0 | 6,423 | 4,277 | 12,482 | 1,316 | 219 |
| 1.1.2 | 144,221 | 70,605 | 9,694 | 18,497 | 430 | 0 | 9,218 | 14,490 | 17,922 | 3,026 | 339 |
| 1.1.3 | 8,118 | 4,087 | 651 | 0 | 0 | 0 | 3,276 | 56 | 0 | 48 | 0 |
| 1.2 | 51,943 | 41,405 | 1,489 | 1,253 | 13 | 0 | 4,366 | 2,838 | 134 | 203 | 242 |
| 1.2.1 | 47,328 | 40,679 | 1,440 | 931 | 13 | 0 | 3,418 | 272 | 134 | 199 | 242 |
| 1.2.2 | 3,640 | 0 | 0 | 322 | 0 | 0 | 753 | 2,565 | 0 | 0 | 0 |
| 2 | 478,070 | 60,098 | 5,620 | 2,838 | 1,874 | 282,500 | 10,859 | 106,403 | 5,891 | 1,761 | 226 |
| 2.1 | 397,819 | 131 | 468 | 1,416 | 1,752 | 282,500 | 7,791 | 97,710 | 5,809 | 59 | 183 |
| 2.1.1 | 92,563 | 5 | 41 | 261 | 0 | 86,195 | 2,078 | 2,301 | 1,645 | 7 | 30 |
| 2.1.2 | 94,929 | 5 | 144 | 228 | 135 | 0 | 918 | 93,494 | 0 | 5 | 0 |
| 2.1.3 | 19,335 | 25 | 108 | 77 | 1,084 | 15,076 | 2,376 | 487 | 65 | 22 | 15 |
| 2.1.4 | 35,981 | 0 | 0 | 0 | 0 | 33,537 | 0 | 13 | 2,370 | 0 | 61 |
| 2.1.5 | 145,791 | 4 | 93 | 139 | 285 | 142,679 | 755 | 131 | 1,664 | 8 | 33 |
| 2.1.6 | 7,552 | 0 | 0 | 711 | 0 | 4,996 | 467 | 1,280 | 65 | 0 | 33 |
| 2.2 | 56,786 | 47,199 | 4,652 | 14 | 4 | 0 | 665 | 2,458 | 82 | 1,669 | 43 |
| 2.3 | 20,129 | 12,767 | 500 | 475 | 118 | 0 | 2 | 6,235 | 0 | 32 | 0 |
| **Total** | **891,045** | **312,334** | **29,452** | **52,691** | **3,154** | **282,500** | **36,767** | **129,911** | **36,429** | **6,733** | **1,074** |

**Table S5.**
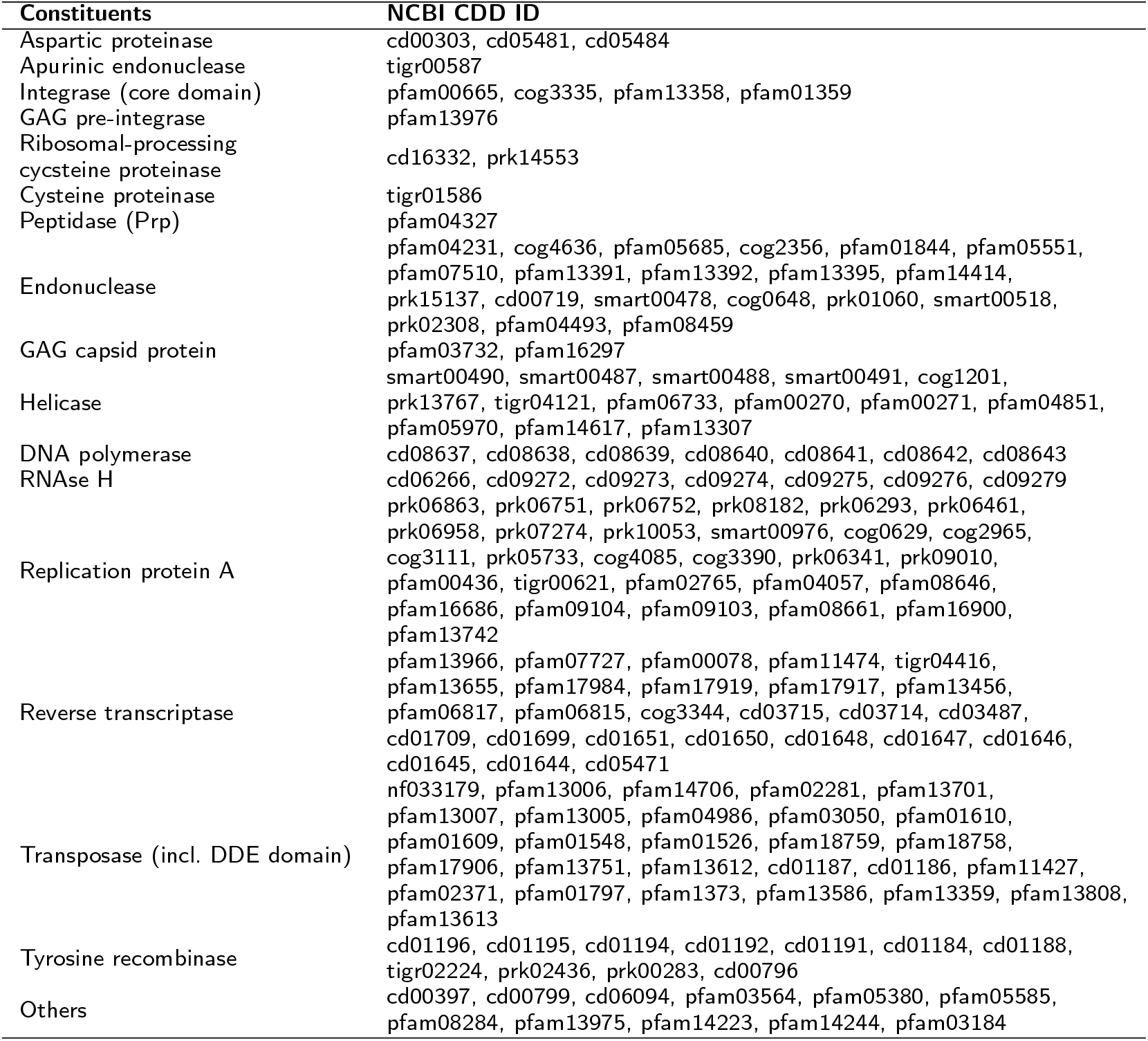
Classification: selection of protein domains. This table lists the selected NCBI CDD PSSM model IDs considered for the protein features used in the classification module.

**Table S6.** Annotation: case study genomes. This diagram shows source, species, sequencing technology, number of sequences, length in bp, strain name and location for the 25 case study genomes.

| Source | Species | Seq. Technology | #Sequences | Length (BP) | Strain name | Strain location |
| --- | --- | --- | --- | --- | --- | --- |
| IRGSP | *Oryza sativa subsp. japo.* | Illumina | 12 | 374,471,240 | Nipponbare | Japan |
| Wormbase WS279 | *Caenorhabditis elegans* | III., PacBio, Nano. | 7 | 102,092,263 | VC2010 | Bristol (UK) |
| Wormbase WS279 | *Caenorhabditis elegans* | Sanger | 7 | 100,286,401 | N2 | Bristol (UK) |
| Wormbase WS279 | *Caenorhabditis elegans* | Illumina | 7 | 98,291,416 | CB4856 | Hawai (USA) |
| Cristian Riccio | *Caenorhabditis elegans* | PacBio, Illumina | 6 | 105,149,162 | SX3383 | Ulupalakua (USA) |
| Cristian Riccio | *Caenorhabditis elegans* | PacBio, Illumina | 6 | 103,998,030 | SX3399 | Wuhan (China) |
| Cristian Riccio | *Caenorhabditis elegans* | PacBio, Illumina | 6 | 104,190,248 | SX3356 | San Francisco (USA) |
| Cristian Riccio | *Caenorhabditis elegans* | PacBio, Illumina | 6 | 103,085,866 | SX3351 | Addisababa (Ethiopia) |
| Cristian Riccio | *Caenorhabditis elegans* | PacBio, Illumina | 6 | 103,400,028 | SX3368 | Adelaide (Australia) |
| Cristian Riccio | *Caenorhabditis elegans* | PacBio, Illumina | 6 | 102,737,178 | SX3287 | Altadena (USA) |
| Cristian Riccio | *Caenorhabditis elegans* | PacBio, Illumina | 6 | 102,467,336 | SX3304 | Amares (Portugal) |
| Cristian Riccio | *Caenorhabditis elegans* | PacBio, Illumina | 6 | 104,412,447 | SX3306 | Auckland (New Zealand) |
| Cristian Riccio | *Caenorhabditis elegans* | PacBio, Illumina | 6 | 102,638,870 | SX3254 | Bristol (UK) |
| Cristian Riccio | *Caenorhabditis elegans* | PacBio, Illumina | 6 | 103,222,210 | SX3292 | Hawai (USA) |
| Cristian Riccio | *Caenorhabditis elegans* | PacBio, Illumina | 6 | 102,808,004 | SX3302 | Hermanville (France) |
| Cristian Riccio | *Caenorhabditis elegans* | PacBio, Illumina | 6 | 102,945,196 | SX3303 | Lake Forest Park (USA) |
| Cristian Riccio | *Caenorhabditis elegans* | PacBio, Illumina | 6 | 103,095,877 | SX3298 | Lisbon (Portugal) |
| Cristian Riccio | *Caenorhabditis elegans* | PacBio, Illumina | 6 | 102,627,405 | SX3295 | Madagascar (Madagascar) |
| Cristian Riccio | *Caenorhabditis elegans* | PacBio, Illumina | 6 | 104,465,563 | SX3367 | Manuka (USA) |
| Cristian Riccio | *Caenorhabditis elegans* | PacBio, Illumina | 6 | 103,159,706 | SX3305 | Palo Alto (USA) |
| Cristian Riccio | *Caenorhabditis elegans* | PacBio, Illumina | 6 | 102,930,261 | SX3296 | Roxel (Germany) |
| Cristian Riccio | *Caenorhabditis elegans* | PacBio, Illumina | 6 | 102,990,752 | SX3286 | Salt Lake City (USA) |
| Cristian Riccio | *Caenorhabditis elegans* | PacBio, Illumina | 6 | 103,010,394 | SX3350 | Southampton (USA) |
| Cristian Riccio | *Caenorhabditis elegans* | PacBio, Illumina | 6 | 103,231,504 | SX3341 | Le Perreux-sur-Marne (France) |
| JST-ACCEL (NIBB) | *Rhizophagus irregularis* | PacBio | 210 | 149,750,837 | DAOM197198 | Quebec (Canada) |

**Table S7.** Classification: benchmark of classifiers. All performance measures reported as average across 10 folds (in %) are supplemented by the standard deviations in brackets (in %). Bold numbers mark the best performance amongst different classifiers within same category. Panel (A) outlines performance measures of the benchmark algorithms reported in their publications (meaning these results were gathered from different datasets and taxonomies, depending on the specific publication). Panel (B) outlines performance measures of several benchmark algorithms to the proposed “RFSB” classifier methodology. All results were calculated based on the same dataset RepBase+PGSB and the same, proposed taxonomy. The measures are reported for taxonomic levels and overall perspective. In addition, the proposed “RFSB” classifier is applied to TransposonDB and reported in the most right column.

| (A) Benchmark of classifiers, performances reported in publications of classifiers (different databases and taxonomies) |  |  |  |  |  |  |  |  |  |
| --- | --- | --- | --- | --- | --- | --- | --- | --- | --- |
|  | Random Even | Random Gaussian | HC_GA | HC_LGA | NLLCPN | TERL | TopDown | RFSB | (TransposonDB) RFSB |
| Overall |  |  |  |  |  |  |  |  |  |
| F1 | 31.03 %<br>(0.32%) | 30.44%<br>(0.26%) | 83.00% | 84.00% | 90.00% | 85.80% | 83.00% | **91.34%**<br>(0.33%) | 89.66%<br>(0.06%) |
| (B) Benchmark of classifiers, performances reproduced on RepBase+PGSB database and proposed taxonomy |  |  |  |  |  |  |  |  |  |
|  | Random Even | Random Gaussian | HC_GA | HC_LGA | NLLCPN | TERL | TopDown | RFSB | (TransposonDB) RFSB |
| Overall |  |  |  |  |  |  |  |  |  |
| ACC | 47.58%<br>(0.27%) | 47.39%<br>(0.38%) | 84.19%<br>(0.41%) | 80.94%<br>(0.37%) | 79.96%<br>(0.28%) | 24.10%<br>(2.91%) | 38.60%<br>(23.40%) | **93.75%**<br>(0.24%) | 93.28%<br>(0.04%) |
| F1 | 31.03 %<br>(0.32%) | 30.44%<br>(0.26%) | 77.91%<br>(0.59%) | 74.02%<br>(0.56%) | 70.84%<br>(0.38%) | 7.11%<br>(0.99%) | 24.43%<br>(24.43%) | **91.34%**<br>(0.33%) | 89.66%<br>(0.06%) |
| MCC | -11.14%<br>(0.54%) | -11.71%<br>(0.63%) | 65.99%<br>(0.86%) | 59.45%<br>(0.79%) | 57.52%<br>(0.60%) | -54.45%<br>(5.41%) | -27.83%<br>(45.19%) | **86.45%**<br>(0.52%) | 84.68%<br>(0.09%) |
| Level 1 |  |  |  |  |  |  |  |  |  |
| ACC | 79.23%<br>(0.09%) | 79.52%<br>(0.07%) | 92.86%<br>(0.13%) | 91.78%<br>(0.27%) | 92.32%<br>(0.10%) | 82.18%<br>(0.54%) | 77.99%<br>(5.75%) | **96.48%**<br>(0.10%) | 96.11%<br>(0.02%) |
| F1 | 29.52%<br>(0.3%) | 29.14%<br>(0.23%) | 75.05%<br>(0.47%) | 71.17%<br>(0.51%) | 70.19%<br>(0.39%) | 0.10%<br>(0.09%) | 23.05%<br>(21.99%) | **88.30%**<br>(0.32%) | 87.74%<br>(0.07%) |
| MCC | 17.35%<br>(0.36%) | 17.18%<br>(0.26%) | 70.92%<br>(0.53%) | 66.44%<br>(0.67%) | 66.70%<br>(0.45%) | -7.36%<br>(0.64%) | 10.31%<br>(25.19%) | **86.27%**<br>(0.38%) | 85.44%<br>(0.09%) |
| Level 2 |  |  |  |  |  |  |  |  |  |
| ACC | 82.60%<br>(0.09%) | 82.91%<br>(0.05%) | 93.63%<br>(0.13%) | 92.72%<br>(0.26%) | 93.26%<br>(0.08%) | 89.21%<br>(0.09%) | 82.33%<br>(3.92%) | **96.99%**<br>(0.09%) | 96.67%<br>(0.02%) |
| F1 | 17.57%<br>(0.38%) | 17.13%<br>(0.23%) | 68.51%<br>(0.73%) | 63.70%<br>(0.70%) | 60.18%<br>(0.43%) | 00.00%<br>(0.00%) | 12.89%<br>(20.23%) | **86.30%**<br>(0.40%) | 85.66%<br>(0.08%) |
| MCC | 7.85%<br>(0.43%) | 7.62%<br>(0.24%) | 65.12%<br>(0.75%) | 59.89%<br>(0.78%) | 59.21%<br>(0.48%) | -1.01%<br>(0.50%) | 3.21%<br>(22.08%) | **84.65%**<br>(0.45%) | 83.81%<br>(0.09%) |
| Level 3 |  |  |  |  |  |  |  |  |  |
| ACC | 86.30%<br>(0.05%) | 86.57%<br>(0.04%) | 92.87%<br>(0.24%) | 91.69%<br>(0.22%) | 90.91%<br>(0.00%) | 90.90%<br>(0.01%) | 86.96%<br>(3.16%) | **97.33%**<br>(0.11%) | 97.76%<br>(0.02%) |
| F1 | 9.63%<br>(0.41%) | 9.32%<br>(0.34%) | 49.93%<br>(2.77%) | 38.67%<br>(4.62%) | 0.00%<br>(0.0%) | 0.00%<br>(0.00%) | 7.96%<br>(12.97%) | **84.95%**<br>(0.59%) | 87.51%<br>(0.10%) |
| MCC | 2.59%<br>(0.42%) | 2.56%<br>(0.34%) | 48.61%<br>(2.44%) | 37.48%<br>(3.24%) | 0.00%<br>(0.00%) | -00.21%<br>(0.17%) | 2.28%<br>(12.87%) | **83.51%**<br>(0.65%) | 86.29%<br>(0.11%) |

**Table S8.**
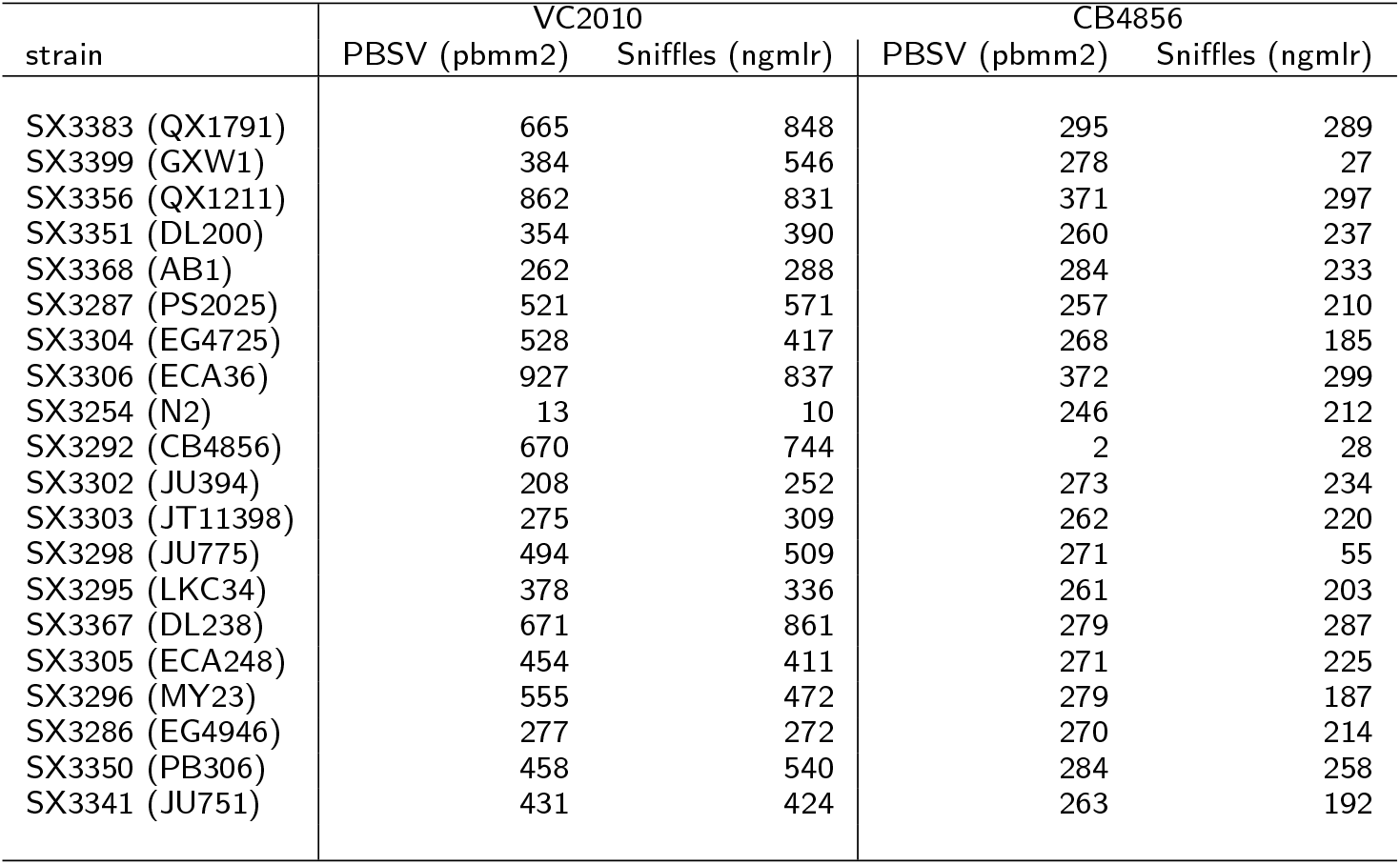
Detection: Number of observed transposition events. This table shows the number of observed transposition events for different probe reference genome combinations, alignment and structural variant calling tools.

| strain | VC2010 |  | CB4856 |  |
| --- | --- | --- | --- | --- |
|  | PBSV (pbmm2) | Sniffles (ngmlr) | PBSV (pbmm2) | Sniffles (ngmlr) |
| SX3383 (QX1791) | 665 | 848 | 295 | 289 |
| SX3399 (GXW1) | 384 | 546 | 278 | 27 |
| SX3356 (QX1211) | 862 | 831 | 371 | 297 |
| SX3351 (DL200) | 354 | 390 | 260 | 237 |
| SX3368 (AB1) | 262 | 288 | 284 | 233 |
| SX3287 (PS2025) | 521 | 571 | 257 | 210 |
| SX3304 (EG4725) | 528 | 417 | 268 | 185 |
| SX3306 (ECA36) | 927 | 837 | 372 | 299 |
| SX3254 (N2) | 13 | 10 | 246 | 212 |
| SX3292 (CB4856) | 670 | 744 | 2 | 28 |
| SX3302 (JU394) | 208 | 252 | 273 | 234 |
| SX3303 (JT11398) | 275 | 309 | 262 | 220 |
| SX3298 (JU775) | 494 | 509 | 271 | 55 |
| SX3295 (LKC34) | 378 | 336 | 261 | 203 |
| SX3367 (DL238) | 671 | 861 | 279 | 287 |
| SX3305 (ECA248) | 454 | 411 | 271 | 225 |
| SX3296 (MY23) | 555 | 472 | 279 | 187 |
| SX3286 (EG4946) | 277 | 272 | 270 | 214 |
| SX3350 (PB306) | 458 | 540 | 284 | 258 |
| SX3341 (JU751) | 431 | 424 | 263 | 192 |

**Table S9.**
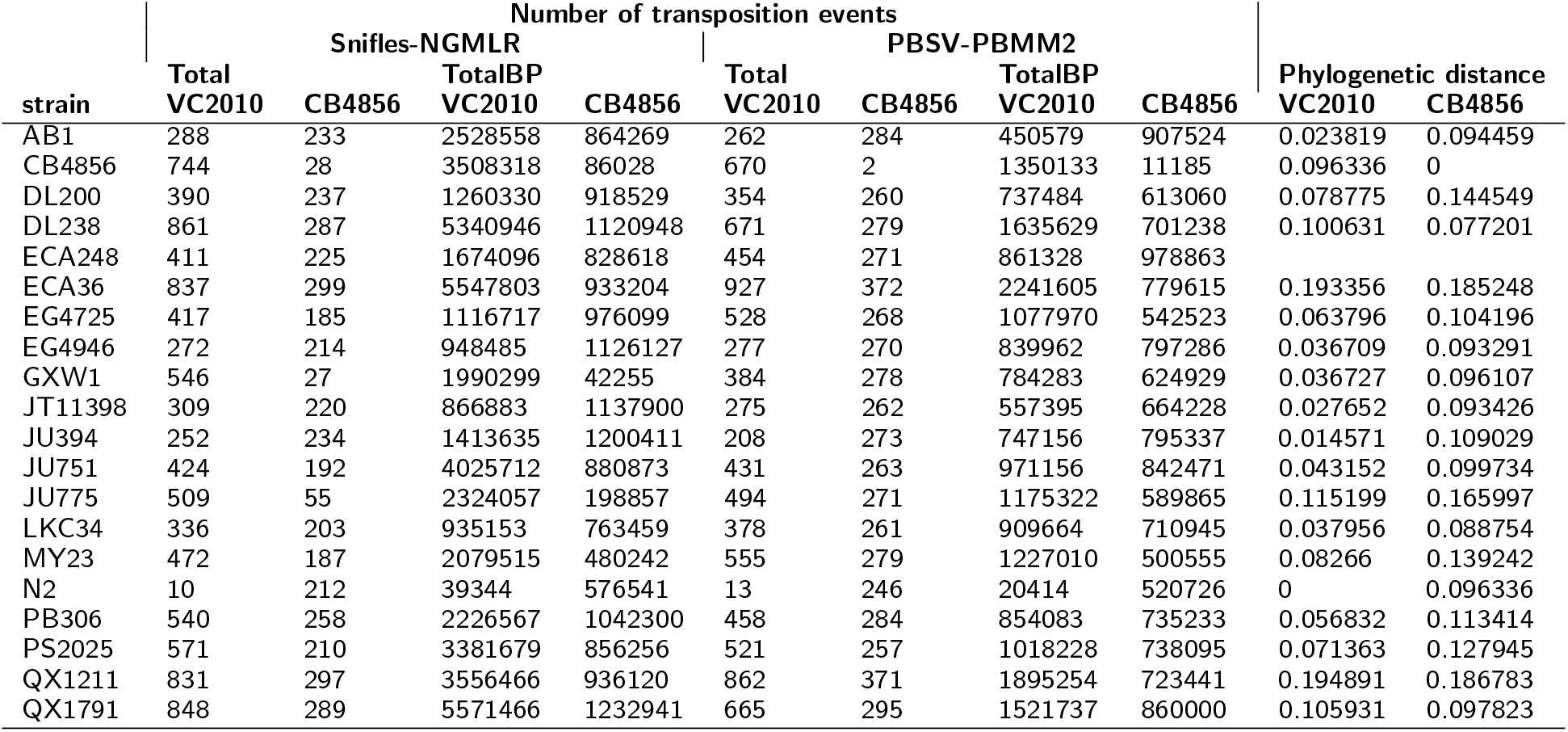
Detection: Genetic distance and number of transposition events found. This table shows the number of observed transposition event candidates, the length of their mask in bp, and the phylogenetic distance of probe and reference genome.

## Footnotes

[1] We use version 23.08 that was the last publicly available version before the paywall was introduced.

